# *ΔG_unfold_ leaderless*, a package for high-throughput analysis of translation initiation regions (TIRs) at the transcriptome scale and for leaderless mRNA optimization

**DOI:** 10.1101/2021.08.27.457836

**Authors:** Mohammed-Husain M. Bharmal, Jared M. Schrader

## Abstract

**Background:** Translation initiation is an essential step for fidelity of gene expression, in which the ribosome must bind to the translation initiation region (TIR) and position the initiator tRNA in the P-site (1). For this to occur correctly, the TIR encompassing the ribosome binding site (RBS) needs to be highly accessible (2-5). ΔG_unfold_ is a metric for computing accessibility of the TIR, but there is no automated way to compute it manually with existing software/tools limiting throughput.

**Results:** *ΔG*_*unfold*_ *leaderless* allows users to automate the ΔG_unfold_ calculation to perform high-throughput analysis. Importantly, *ΔG*_*unfold*_ *leaderless* allows calculation of TIRs of both leadered mRNAs and leaderless mRNAs which lack a 5’ UTR and which are abundant in bacterial, archaeal, and mitochondrial transcriptomes (4, 6, 7). The ability to analyze leaderless mRNAs also allows one additional feature where users can computationally optimize leaderless mRNA TIRs to maximize their gene expression (8, 9).

**Conclusions:** The ΔG_unfold_ leaderless package facilitates high-throughput calculations of TIR accessibility, is designed to calculate TIR accessibility for leadered and leaderless mRNA TIRs which are abundant in bacterial/archaeal/organellar transcriptomes and allows optimization of leaderless mRNA TIRs for biotechnology.

## Background

ΔG_unfold_ is a metric to compute accessibility of translation initiation region (TIR) which is an important determinant of start codon selection (10). Translation initiation is an essential step for fidelity of gene expression, in which the Ribosome must bind to the TIR and position the start codon in the P-site with the initiator tRNA (1). For this to occur, the translation initiation region (TIR) encompassing the ribosome binding site (RBS) needs to be highly accessible, lacking secondary structure (2-5). In contrast to calculating the minimum free energy (mfe) of the TIR region using nearest neighbor approaches, ΔG_unfold_ provides information about the TIR region boundary by providing the site of ribosome binding, giving an accessibility measure that better correlates with translation efficiency (11-14). ΔG_unfold_ as the metric for computing accessibility has gained appreciation but there is no automated way to compute it with existing softwares/tools. This currently limits the use of ΔG_unfold_ as the low throughput of calculation has made it difficult to perform transcriptome level analyses. Importantly, there are softwares and algorithms available which can predict secondary structure of any mRNA sequence and calculate the minimum free energy(mfe) of the resulting secondary structure. With these tools, one can in principle calculate the ΔG_unfold_ of any structure, however, this requires multiple steps to be executed sequentially and the constrained structure needs to be generated and inputted manually based on the start codon position and TIR location. So, though computation of the ΔG_unfold_ for a single sequence is possible, it is not a practical solution for the entire transcriptome or multiple sequences. Therefore, this *ΔG*_*unfold*_ *leaderless* software will make high-throughput transcriptome-level computation of TIR accessibility feasible.

Importantly, *ΔG*_*unfold*_ *leaderless* also has TIR optimization program which can be used to optimize expression of proteins in bacteria with predominant non-SD translation initiation mechanism, unlike *E*.*coli. E. coli* has been widely used for expression of recombinant proteins as it is very well characterized and various genetic tools are readily available (15). However, two key challenges for heterologous protein expressions are efficient synthesis (dictated by rates of transcription and translation) and misfolding. Therefore, *E. coli* system cannot be the best choice for all the applications and other hosts are required influenced by the application and availability of tool for their genetic manipulation (15). For instance, translation initiation in *C. crescentus* is not predominantly Shine-Dalgarno (SD) dependent unlike *E. coli*, and only approximately 23% of genes initiate translation via SD mediated mechanism (16-18) with some organisms containing SD sites in as few as 8% (7, 19). Further, RNA-seq based transcription mapping experiments have found that many bacterial mRNAs are “leaderless” and begin directly at the AUG start codon (20-22), and that this type of mRNAs are abundant in pathogens such as *M. tuberculosis* and in the mammalian mitochondria (6). In C. crescentus approximately 17% of its transcriptome is leaderless (16). For leaderless mRNAs, three factors are known to affect translation initiation efficiency: accessibility of TIRs (ΔG_unfold_), start codon identity and leader length with higher accessibility of TIR, AUG as start codon and any absence of short leaders are most efficient (7, 23-30). Therefore, these properties can be exploited to optimize expression of proteins in such organisms having predominantly leaderless translation initiation machinery. A similar approach has been used in *E*.*coli*, in which synonymous mutations were rendered in 5’ coding region to elevate the expression of the reporter protein (8). Using this part of the *ΔG*_*unfold*_ *leaderless* package, the user can input the coding sequence (CDS) and then the program will alter the start codon to AUG if there is any other codon and make synonymous mutations in the TIR and compute ΔG_unfold_ for all possible combinations and output most preferred sequence in terms of least ΔG_unfold_ or highest accessibility.

## Implementation

### General requirements

ΔG_unfold_ leaderless is an open-source software available as a linux shell script available through github (https://github.com/schraderlab/dGunfold_program) based upon the RNAfold and dot2ct packages (31, 32). ΔG_unfold_ represents the energy required to unfold the mRNA’s ribosome binding site (RBS) (Fig 1A) (10) which can be useful in studies of translation initiation. There are 3 subprograms available based on the need of the user, which are for transcriptome studies, leaderless mRNA TIR optimization, or calculation of custom sequences. Before running ΔG_unfold_ leaderless, the user also needs to install RNAfold (32) and RNAstructure (31) programs as described in the README file. The computational pipeline of the three subprograms are described in the following sections, and detailed instructions to run the software on linux can be found in the README file.

**Figure 1:**
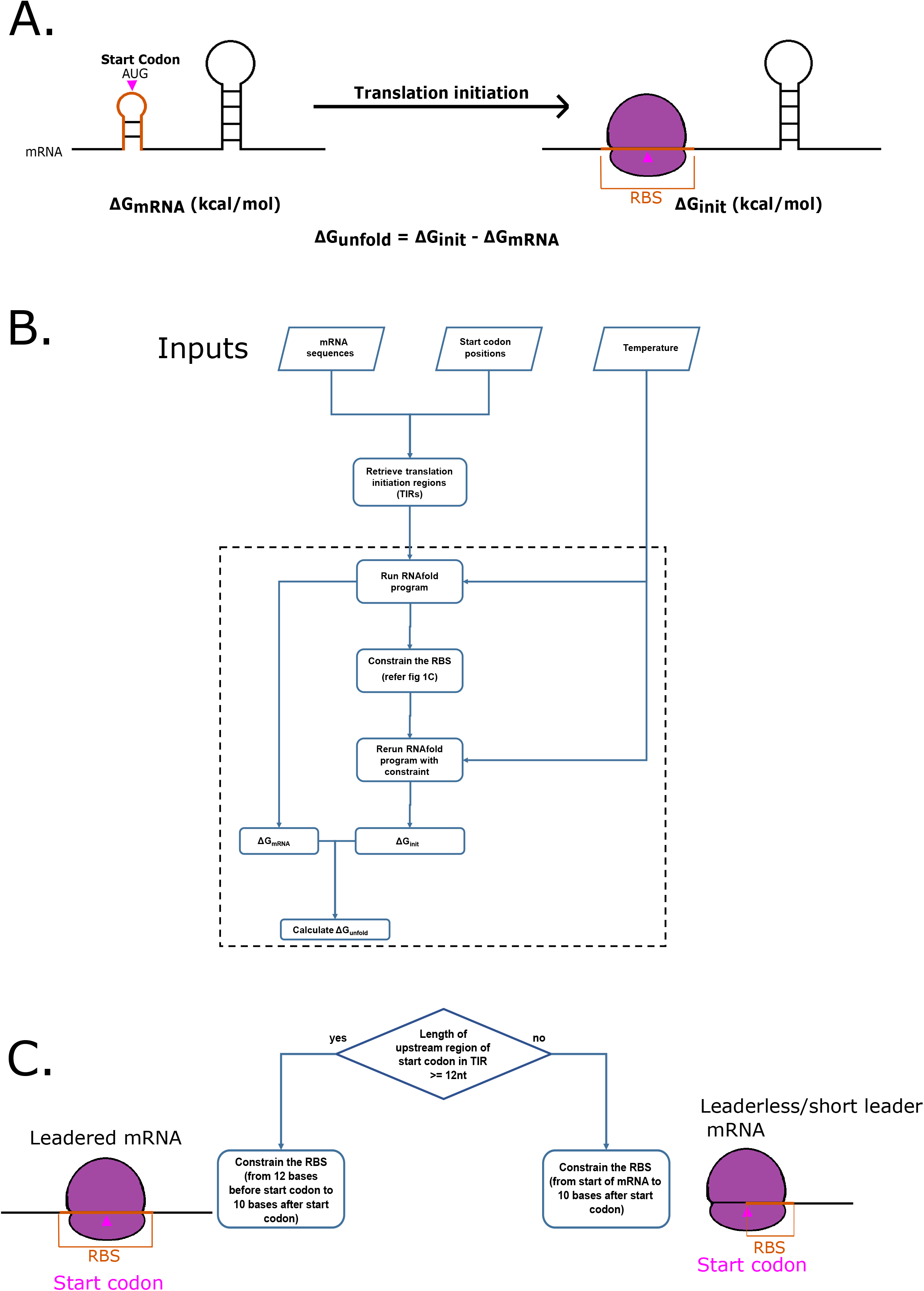
ΔG_unfold_ core computation. A) Graphical representation showing the unfolding of the ribosome binding site (RBS) in the translation initiation region (TIR), facilitating binding of the ribosome to the RBS. Also, showing the calculation for ΔG_unfold_. B) Flowchart of execution of ΔG_unfold_ leaderless program with user input sequences. The core component of this pipeline is shown within the dotted box. An example output file is included in Table 1. C) Algorithm for accounting for short/leaderless mRNAs. Flowchart showing RBS length selection for constraining based on the length of 5’ UTR.

### The basic ΔG_unfold_ Computational Pipeline

The core section of this program is ΔG_unfold_ calculation (Fig 1). In order to calculate the ΔG_unfold_ of multiple RNAs, a .txt file with all the RNA sequences and a second .txt file with start codon positions in the same order as the sequences are inputted into the program (Fig 1B). The user can also define the temperature in which they want to calculate ΔG_unfold_. The ΔG_unfold_ leaderless package will then calculate the minimum free energy of the mRNA structure using RNAfold (ΔG_mRNA_), and will then constrain the ribosome binding site (RBS) to be single stranded as an approximation for ribosome accessibility, and then calculate the minimum free energy of the constrained mRNA structure (ΔG_init_) (Fig 1B). With these two measures, the ΔG_unfold_ can be calculated for each RNA (Fig 1B). The RBS region is defined by the size of a ribosome footprint surrounding the start codon (25nt) which is dynamically adapted to smaller sizes for mRNAs with short 5’ UTRs, or leaderless mRNAs which completely lack 5’ UTRs (Fig 1C). To constrain the structure as single-stranded, all base-pairs within the RBS are constrained to be single stranded before recalculating the ΔG_init_. The output file contains six columns: RNA sequence, dot-bracket mfe structure, constrained structure, ΔG_mRNA_ values, ΔG_init_ values and ΔG_unfold_ values (Table 1). Users are allowed to additionally control the size of the TIR region for analysis (default = 50nt) which provide additional flexibility to the user.

### The ΔG_unfold_ pipeline for transcriptome scale analysis

Bacterial transcriptome architecture maps provide an ideal platform to analyze TIR accessibility on a transcriptome-wide scale. Notably, transcription start site (TSS) data and RNA-seq density data have been used previously to experimentally map bacterial transcriptomes (33). If such TSS data are available, the user can input this together with the genbank file of the respective genome containing the CDS annotations and an operon map to build a transcript architecture model that allows the best starting point for transcriptome analyses. TSSs are assigned to transcripts by searching within 300nts upstream of the start codon of either the CDS (or the leading CDS in an operon). Multiple TSS sites may be assigned to a single operon/gene and in such cases the different mRNA isoforms are included in the transcript architecture map. Once the TSS sites are defined to each operon, the ΔG_unfold_ is calculated for the TIR of each CDS in each mRNA transcriptional units. If no TSS data is provided, the software can approximate each CDS as having an mRNA leader of 25 nts, although this leads to uncertainly about short leadered and leaderless mRNAs. Overall, the ΔG_unfold_ leaderless transcriptome analysis runs within 30 min on a budget desktop cpu for a dataset equivalent to the largest known bacterial transcriptome, suggesting that this software can be run to study any bacterial transcriptome (Fig 2B).

**Figure 2:**
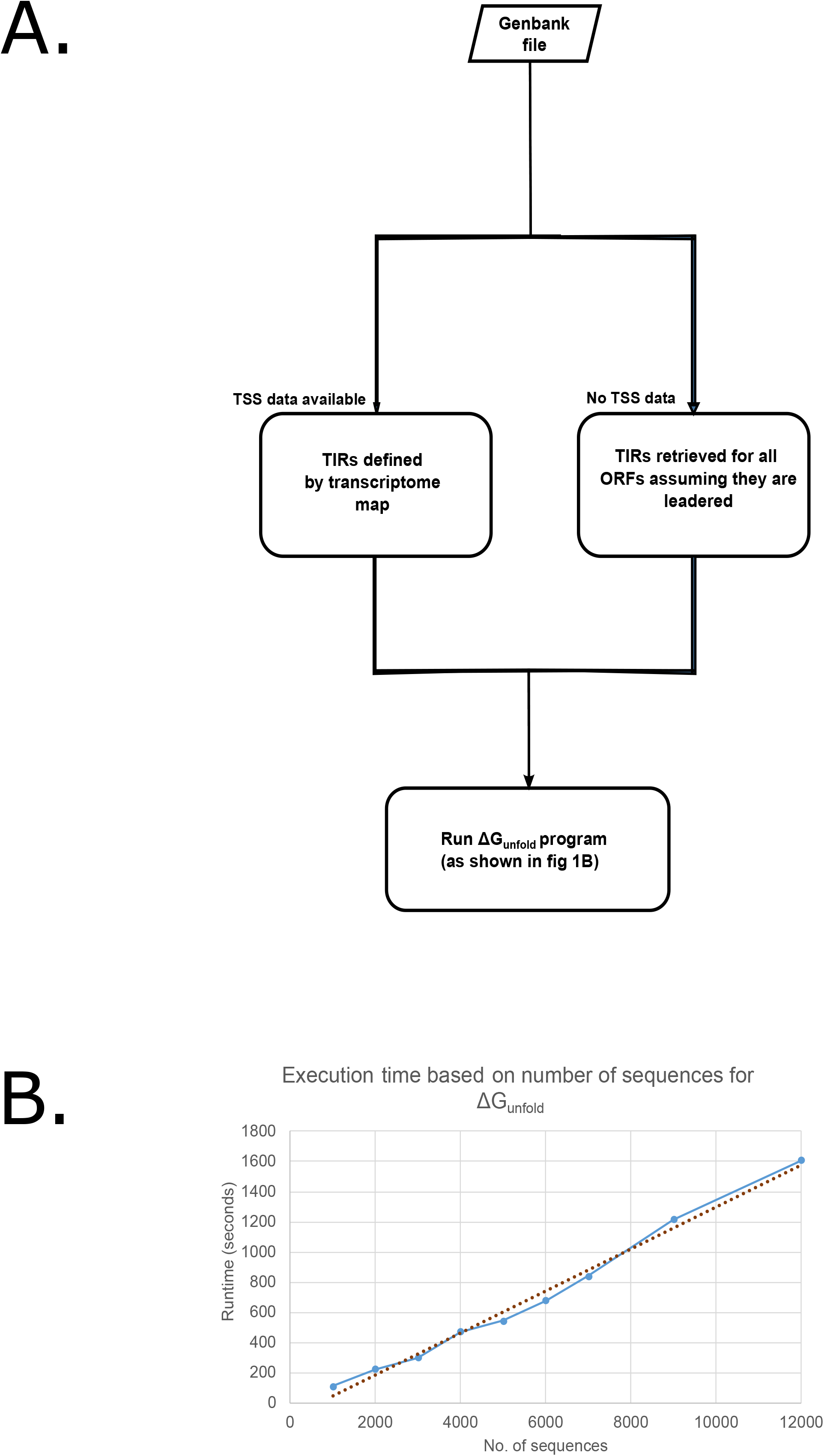
Transcriptome-wide ΔG_unfold_ computation. A) Flowchart of execution of ΔG_unfold_ leaderless program for the entire transcriptome. If TSS data is available, the exact transcriptional units defined by transcription start site data can be used (left), or if not available, each transcriptional unit will be calculated as if it contained a leader of >=25 nts. An example output file is included in Table 2. B) Scatter plot showing the time required to execute the core component of the program (Y-axis) for different number of sequences (X-axis). The maximum size of the X-axis was chosen to be equivalent to the largest bacterial genome identified. Analysis was performed using an AMD Ryzen 3200g desktop cpu.

### ΔG_*unfold*_ for optimizing leaderless mRNA translation initiation

The lack of secondary structure is a strong determinant of the translation initiation efficiency of leaderless mRNAs (9, 34, 35). As some organisms with abundant leaderless mRNAs are important for biotechnology, we adapted *ΔG*_*unfold*_ *leaderless* to generate mutant versions with higher accessibility to optimize recombinant gene expression (Fig 3). The user inputs the sequence of their leaderless mRNA to optimize, and all possible synonymous codon mutations in the RBS are generated and output to the user in a rank ordered list from lowest to highest *ΔG*_*unfold*_. The user can then select those variants with optimal properties for optimal expression testing.

**Figure 3:**
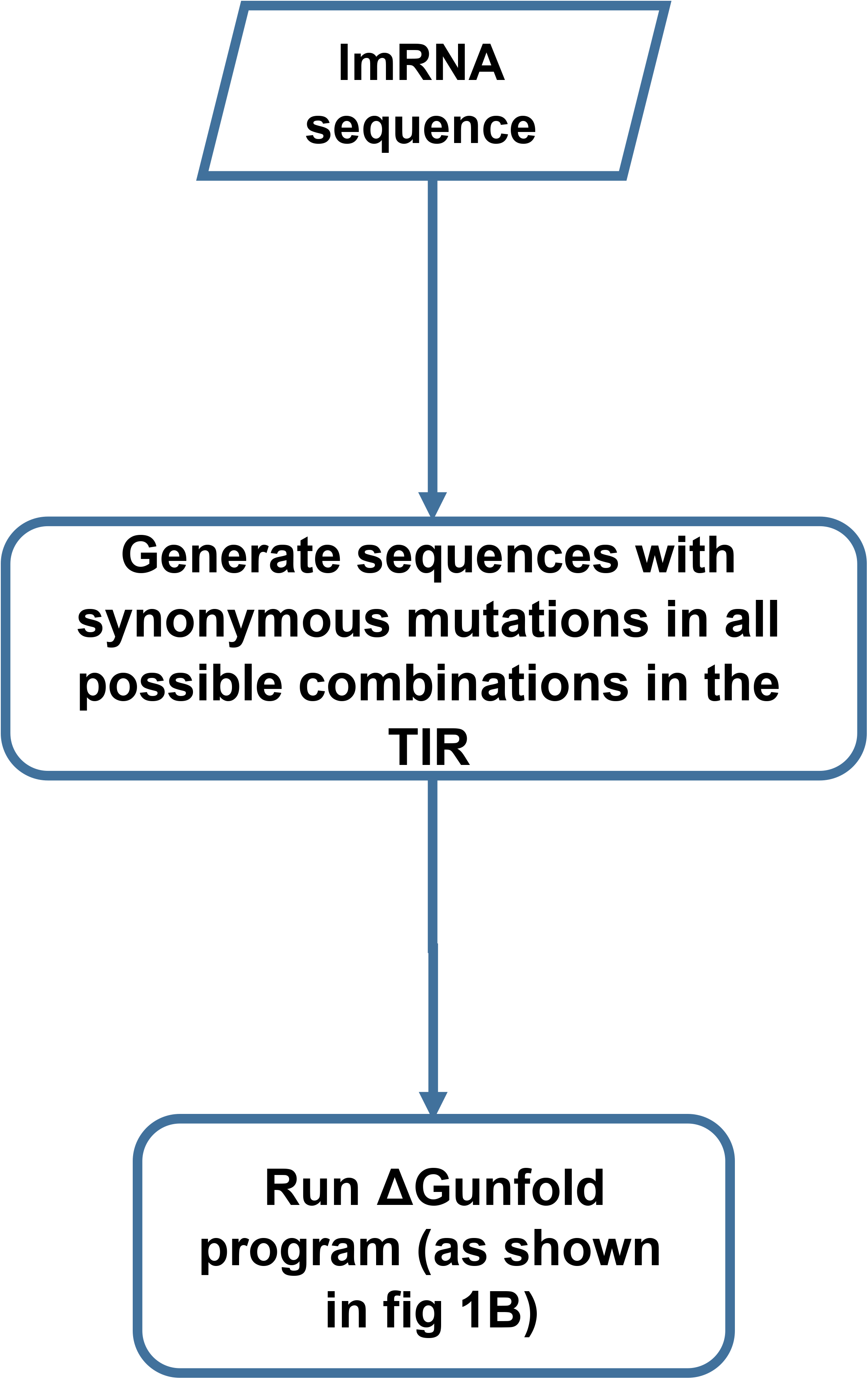
Translation initiation region (TIR) accessibility optimization program. Flowchart of execution of ΔG_unfold_ leaderless program for the TIR accessibility optimization. Optimization is performed using synonymous codon mutations in the RBS and providing the user a rank-order list of all variants based on their ΔG_unfold_ (Table 3).

## Conclusions

*ΔG*_*unfold*_ *leaderless* automates the computation of mRNA TIR accessibility for large number of sequences allowing transcriptome-level analysis. Indeed, runtimes allow users to quickly calculate *ΔG*_*unfold*_ across a whole transcriptome using a budget desktop computer (Figure 2B). In addition, *ΔG*_*unfold*_ *leaderless* can also be used for biotechnology purposes to help optimize the translation initiation of recombinantly expressed leaderless mRNAs for biotechnology purposes. Together, these functions make *ΔG*_*unfold*_ *leaderless* useful for the study of TIR accessibility on mRNA translation.

## Availability and requirements

**Project name:** ΔG_unfold_ leaderless

**Project home page:** https://github.com/schraderlab/dGunfold_program

**Operating system(s):** Linux/Unix command line

**Programming language:** Python/shell script

**Other requirements:** Python 3.0 or higher, Biopython

**License:** GNU General Public License v3.0

**Any restrictions to use by non-academics:** e.g. licence needed

## List of abbreviations

TIR: translation initiation region
RBS: ribosome binding site
Mfe: minimum free energy
SD: Shine-Dalgarno

## Declarations

### Ethics approval and consent to participate

“Not applicable”

### Consent for publication

“Not applicable”

### Availability of data and materials

All relevant python files and example datasets are freely available in the ΔGunfold leaderless github repository https://github.com/schraderlab/dGunfold_program

### Authors’ contributions

MHB wrote software. MHB and JMS tested software and wrote manuscript.

## Acknowledgements

The authors thank members of the Schrader lab for critical feedback. The authors thank James Aretakis and Aishwarya Ghosh for beta-testing and feedback on usage.

## Table and Figure legends

Table 1: Output file format of ΔG_unfold_ leaderless program with user input sequences.

**(XLSX)**.

Table 2: Output file formats of ΔG_unfold_ leaderless program for transcriptome analysis.

**(XLSX)**.

Table 3: Output file format of ΔG_unfold_ leaderless program for TIR accessibility optimization.

**(XLSX)**.

